# Culture under normoxic conditions and enhanced virulence of phase II *Coxiella burnetii* transformed with a RSF1010-based shuttle vector

**DOI:** 10.1101/747220

**Authors:** Shengdong Luo, Zemin He, Zhihui Sun, Yonghui Yu, Yongqiang Jiang, Yigang Tong, Lihua Song

## Abstract

*Coxiella burnetii* is a Gram-negative, facultative intracellular microorganism that can cause acute or chronic Q fever in human. It was recognized as an obligate intracellular organism until the revolutionary design of an axenic cystine culture medium (ACCM). Present axenic culture of *C. burnetii* strictly requires a hypoxic condition (<10% oxygen). Here we investigated the normoxic growth of *C. burnetii* strains in ACCM-2 with or without tryptophan supplementation. Three *C. burnetii* strains - Henzerling phase I, Nine Mile phase II and a Nine Mile phase II transformant, were included. The transformant contains a pMMGK plasmid that is composed of a RSF1010 ori, a repABC operon, an eGFP gene and a kanamycin resistance cassette. We found that, under normoxia if staring from an appropriate concentration of fresh age inocula, Nine Mile phase II can grow significantly in ACCM-2 with tryptophan, while the transformant can grow robustly in ACCM-2 with or without tryptophan. In contrast, long-term frozen stocks of phase II and its transformant, and Henzerling phase I of different ages had no growth capability under normoxia under any circumstances. Furthermore, frozen stocks of the transformant consistently caused large splenomegaly in SCID mice, while wild type Nine Mile phase II induced a lesser extent of splenomegaly. Taken together, our data show that normoxic cultivation of phase II *C. burnetii* can be achieved under certain conditions. Our data suggests that tryptophan and an unknown temperature sensitive signal are involved in the expression of genes for normoxic growth regulated by quorum sensing in *C. burnetii*.

## Introduction

*Coxiella burnetii* is a Gram-negative, facultative intracellular pathogen and the causative agent of human Q fever. The primary transmission route for human is through inhalation of contaminated aerosols from the secretions and excretions of domestic ruminants (1). *C. burnetii* infections in humans may manifest as an acute disease (mainly as a self-limiting febrile illness, pneumonia, or hepatitis) or as a chronic disease (mainly endocarditis in patients with previous valvulopathy) (2). The majority (∼50-60%) of human infections are asymptomatic (2, 3). Chronic infections are rare but can be fatal if untreated. *C. burnetii* is a significant cause of culture-negative endocarditis in the United States (4). Treatment of chronic infections is challenging and currently requires a combined antibiotic therapy with doxycycline and hydroxychloroquine for at least 18 months (5). Worldwide only one vaccine for Q fever called Q-Vax is licensed in Australia to protect high risk populations (6).

*C. burnetii* has two phase variants. Virulent *C. burnetii* isolated from natural sources and infections is defined as phase I. It produces full-length LPS that may play an important role in *C. burnetii* persistent infections by masking toll-like receptor ligands from innate immune recognition by human dendritic cells (7). LPS from phase I *C. burnetii* contains two unique biomarkers of methylated sugars (virenose and dihydrohydroxystreptose) at its O-specific chain. When extensively passaged in immunoincompetent hosts, virulent phase I *C. burnetii* mutates to avirulent phase II (8, 9). LPS from phase II *C. burnetii* is severely truncated and only contains lipid A and partial core oligosaccharide. Lipid A, the basal component of LPS, is essential for *C. burnetii* growth in macrophage-like THP-1 cells but nonessential in non-phagocytic cells (10).

The recent description of an axenic culture medium and the subsequent modified axenic culture media provide invaluable tools for *C. burnetii* research (11-15). The first generation of axenic culture of *C. burnetii* requires a complex nutrient-rich medium (Acidified Citrate Cysteine Medium, ACCM) and microaerophilic conditions with 2.5% oxygen and 5% carbon dioxide (11). A second-generation medium (ACCM-2) was generated by replacing fetal bovine serum with methyl-beta-cyclodextrin (12), then was improved by supplementing tryptophan (14, 15). A nutritionally defined medium (ACCM-D) further improved *C. burnetii* growth especially with increased bacterial viability (13). The continuous improvement of axenic culture systems significantly facilitates the development of the *C. burnetii* field.

Phase I *C. burnetii* is a category B select agent with potential for illegitimate use and requires a biosafety level-3 laboratory for culture (16). Phase II *C. burnetii* is approved for use at biosafety level-2 and is widely used for studying the biology and pathogenesis in tissue culture. The caveat of working with phase II *C. burnetii* is the lack of animal models for assessing *in vivo* pathogenesis. To promote the identification of virulence factors in phase II *C. burnetii*, an SCID mouse model was recently established (17). An alternative infection model using larvae of the greater wax moth *Galleria mellonella* was also used to identify virulence factors in phase II *C. burnetii* (18, 19).

In this study, we characterized the growth of various *C. burnetii* strains under different oxygen concentrations in ACCM-2 with or without tryptophan supplementation. We found that, under normoxia if starting from an appropriated concentration of freshly age inocula, *C. burnetii* Nine Mile phase II had significant propagation in ACCM-2 supplemented with tryptophan. Under same conditions, a Nine phase II transformant that contains a RSF1010-based shuttle vector can have robust growth in ACCM-2 with or without tryptophan. Phase I *C. burnetii* had no apparent growth under normoxia in ACCM-2 with or without tryptophan. Long-term frozen stocks of all tested strains unanimously failed to have normoxic growth. In the SCID mouse infection model, compared to wild type Nine Mile phase II, frozen stocks of its transformant caused larger splenomegaly. Our data suggest that *C. burnetii* retains functional genes for aerobic growth, and the expression of these genes might be regulated by a temperature sensitive signal of the quorum sensing system. Our data also suggest that the expression of genes relating to aerobic growth is associated with enhanced *in vivo* virulence in *C. burnetii*.

## Materials and Methods

### *C. burnetii* strains

*C. burnetii* Nine Mile phase II and Henzerling phase I are from our laboratory strain collection. *C. burnetii* transformant -NMII*pMMGK* was constructed by transforming Nine Mile phase II with a shuttle vector pMMGK by electroporation (20). The pMMGK plasmid is composed of a RSF1010 ori, a repABC operon, an eGFP gene and a kanamycin resistance cassette. Construction of pMMGK was described previously (10).

### Cultivation of *C. burnetii*

Seed stocks of *C. burnetii* Nine Mile phase II, Henzerling phase I and phase II transformant NMII*pMMGK* were collected from ACCM-2 cultures at 37°C under 2.5% oxygen, and were stored in fresh ACCM-2 at -80°C. Different ages and concentrations of inocula were used to characterize *C. burnetii* growth in two media (ACCM-2 with or without tryptophan supplementation) under hypoxia or normoxia. Inocula were added in 150 μL media per well in 96 well plates (Corning). The plates were placed in a tri-gas CO_2_ incubator (2.5% oxygen and 5% CO_2_) or a CO_2_ incubator (5% CO_2_) for seven days at 37°C. Each culture condition has six replicates.

### Quantitative PCR

Genome equivalent (GE) of *C. burnetii* was quantified by using qPCR as previously described with minor modifications (21). Bacterial bodies of *C. burnetii* were collected by centrifugation at 15,000 ×g for 30 min. Total DNA was extracted with TIANamp N96 Blood DNA Kit (Tiangen). Genome copy numbers were determined by Taqman probe qPCR specific to *dotA* by using a ViiA™ 7 Real-Time PCR System (Applied Biosystems). PCR conditions were as follows: initial denaturation at 94°C for 10 min, followed by 40 cycles of amplification at 94°C for 15 s, 60°C for 1 min.

### Mouse infection

Female SCID (CB17/Icr-*Prkdc*^*scid*^/IcrlcoCrl) mice were purchased from Vital River Laboratory Animal Technology Co., Ltd (Beijing, China). Two separate mouse infection experiments were conducted. Mice were infected with mock or 1×10^8^ GE *C. burnetii* in 200 μL PBS by intra-peritoneal injection. In the first experiment, 18 four weeks old mice were divided into three groups: control (PBS) group and two infection groups. Fresh age (less than one day old) inocula of Nine Mile phase II and NMII*pMMGK* were used. In the second experiment, 9 five weeks old mice were averaged into three infection groups: one month old Nine Mile phase II, one-month old NMII*pMMGK* and 19 months old NMII*pMMGK*. At indicated days post-infection, mice were weighted and sacrificed to harvest spleens to determine splenomegaly (spleen weight/body weight). Each spleen was homogenized in 2 mL PBS. Total DNAs from 20 μL of each tissue homogenate were purified with DNeasy Blood & Tissue Kit (Qiagen). *C. burnetii* GEs in spleens were determined by quantitative PCR as described above. All animal procedures were carried out in strict accordance with the guidelines for the Care and Use of Laboratory Animals of the National Ministry of Health of China.

### Statistical Analysis

A two-tailed Student *t* test was used for qPCR analysis of *C. burnetii* growth yields under various conditions and was used for comparison of splenomegaly and spleen bacterial loads of different infection groups.

## Results

### Phase II *C. burnetii* transformants can grow robustly under normoxia in ACCM-2 with or without tryptophan supplementation

Current axenic culture of *C. burnetii* requires a microaerophilic environment (2.5% oxygen) (14). But interestingly, *C. burnetii* encodes terminal oxidases-cytochrome *o* (encoded by *cyoABCDE*) and cytochrome *bd* (encoded by *cydAB*), which are typically associated with aerobic and microaerophilic respiration, respectively (22). The presence of cytochrome *o* suggests the possibility of *C. burnetii* replication under aerobic conditions. In our attempts to culture *C. burnetii* in regular CO_2_ incubators (∼20% oxygen), we found that a *C. burnetii* Nine Mile phase II transformant-NMII*pMMGK* can grow robustly (∼3 logs increase) if starting from a reasonably high inoculum concentration in the regular ACCM-2 medium (without tryptophan supplementation) (Figure 1A). The pMMGK plasmid is composed of the pMMB207 backbone (a RSF1010 *ori* and a repABC operon), an eGFP gene and a kanamycin resistance cassette (10). This initial finding of *C. burnetii* normoxic growth was made in T-flask cultures. To thoroughly characterize the influence of multiple culture parameters on *C. burnetii* growth, 96 well plates were used for *C. burnetii* cultivation. Tryptophan supplementation can improve the viability of *C. burnetii* in ACCM-2 (14). When cultured in ACCM-2 with tryptophan supplementation, NMII*pMMGK* displayed similar growth characteristics, though there was a trend of a bigger yield in ACCM-2 with tryptophan supplementation (Figure 1B).

**Figure 1.**
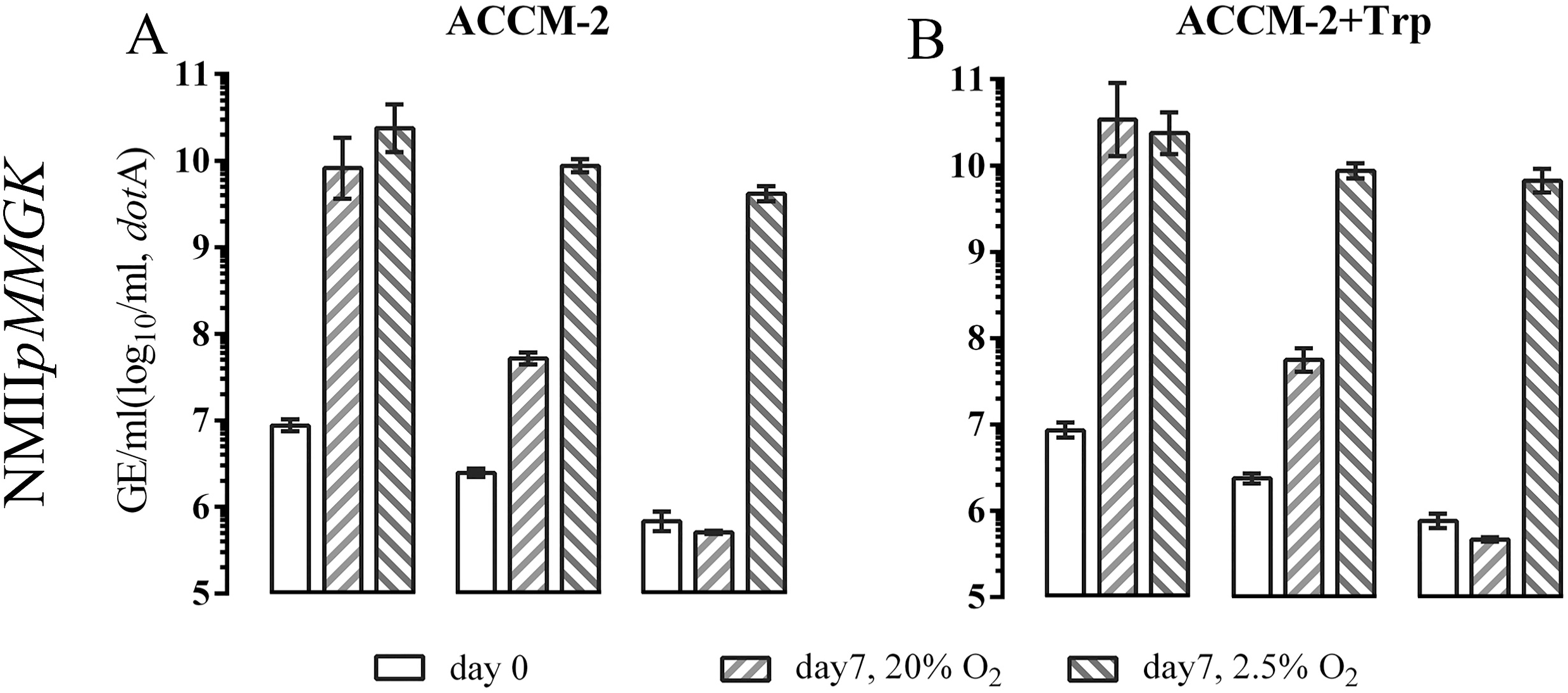
Growth of *C. burnetii* NMII*pMMGK* in ACCM-2 **(A)** and ACCM-2 with tryptophan supplementation **(B)** at 2.5% and 20% oxygen. At 2.5% oxygen, regardless of its inoculum concentration, the NMII*pMMGK* strain consistently has robust growth in ACCM-2 and ACCM-2 plus tryptophan. At 20% oxygen, in both ACCM-2 and ACCM-2 plus tryptophan, the growth yield of NMII*pMMGK* ranges from 0-3.6 logs, depending on its inoculum concentration.

### Tryptophan improved axenic growth of phase II *C. burnetii* under normoxia

The axenic growth of wild type phase I and phase II *C. burnetii* in early generations of ACCMs without tryptophan supplementation has been extensively investigated and was reported to be microaerophilic (2.5% oxygen) (11, 12). We repeated the cultivation of wild type phase I and phase II *C. burnetii* in ACCM-2 without tryptophan at different oxygen concentrations (Figure 2AC). In ACCM-2 without tryptophan supplementation, these two strains grew normally at 2.5% oxygen but had no significant growth at 20% oxygen, which is consistent with previous reports. In ACCM-2 with tryptophan at 2.5% oxygen, both strains had similar growth yields (Figure 2BD). Interestingly, in ACCM-2 with tryptophan at 20% oxygen, wild type phase II *C. burnetii* grew significantly (∼2.5 logs increase) if starting from a high concentration of fresh age inocula (∼4×10^6 GE/mL), while phase I *C. burnetii* consistently had no obvious growth. These results show tryptophan has subtly different effects on the axenic growth of wild type phase I and phase II strains.

**Figure 2.**
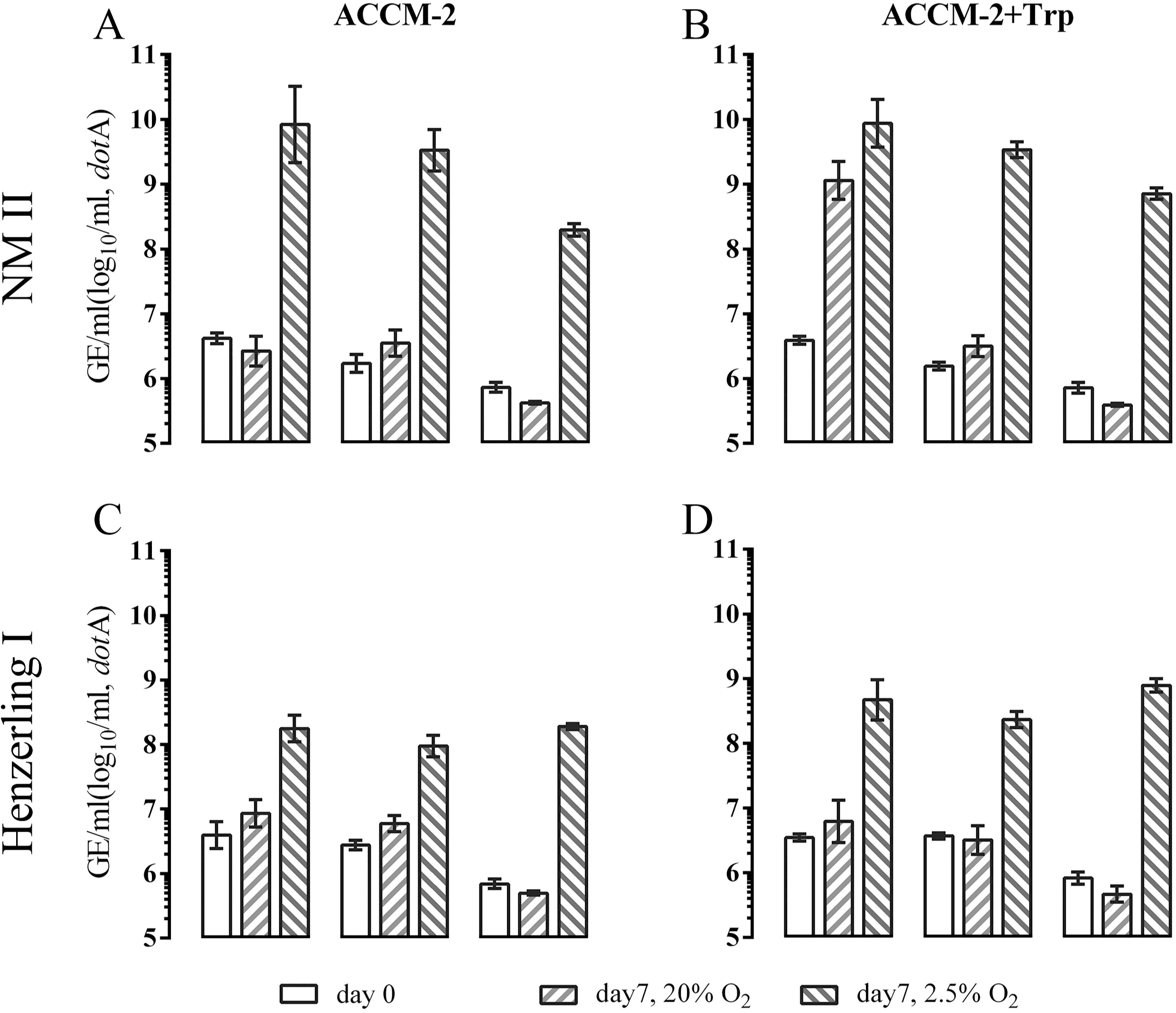
Growth of wild type *C. burnetii* phase I and phase II strains in ACCM-2 and ACCM-2+Trp at 2.5% and 20% oxygen. In ACCM-2, at 20% oxygen, the wild type NM II and Henzerling I strains have no significant growth. In ACCM-2 plus tryptophan, at 20% oxygen, the growth of NM II ranges from 0-2.5 logs, depending on its inoculum concentration, while the Henzerling I strain consistently fails to grow.

### Long-term frozen stocks of *C. burnetii* strains failed to have apparent propagation under normoxia

The cultivation of NMII*pMMGK* at 20% oxygen was constantly repeated. Its viability at 2.5% oxygen in ACCM-2 with or without tryptophan had no obvious variation after long-term frozen (>three months) in -80°C (Figure 3AB). Interestingly, its growth ability at 20% oxygen in either medium was completely abolished after long-term frozen. Similar with NMII*pMMGK*, wild type Nine Mile phase II also failed to replicate at 20% oxygen in ACCM-2 with tryptophan (Figure 3CD). The growth deficiency of wild type Henzerling phase I at 20% oxygen in either medium was consistent (Figure 3EF). Compared to growth of freshly cultivated inocula (Figure 2), lower fold change of growth yield of long-term frozen inocula was observed in either medium (Figure 3C-F), suggesting low viability of both wild type frozen stocks. Taken together, our results suggest that a temperature sensitive factor is associated with the expression of genes involved in *C. burnetii* aerobic growth and viability.

**Figure 3.**
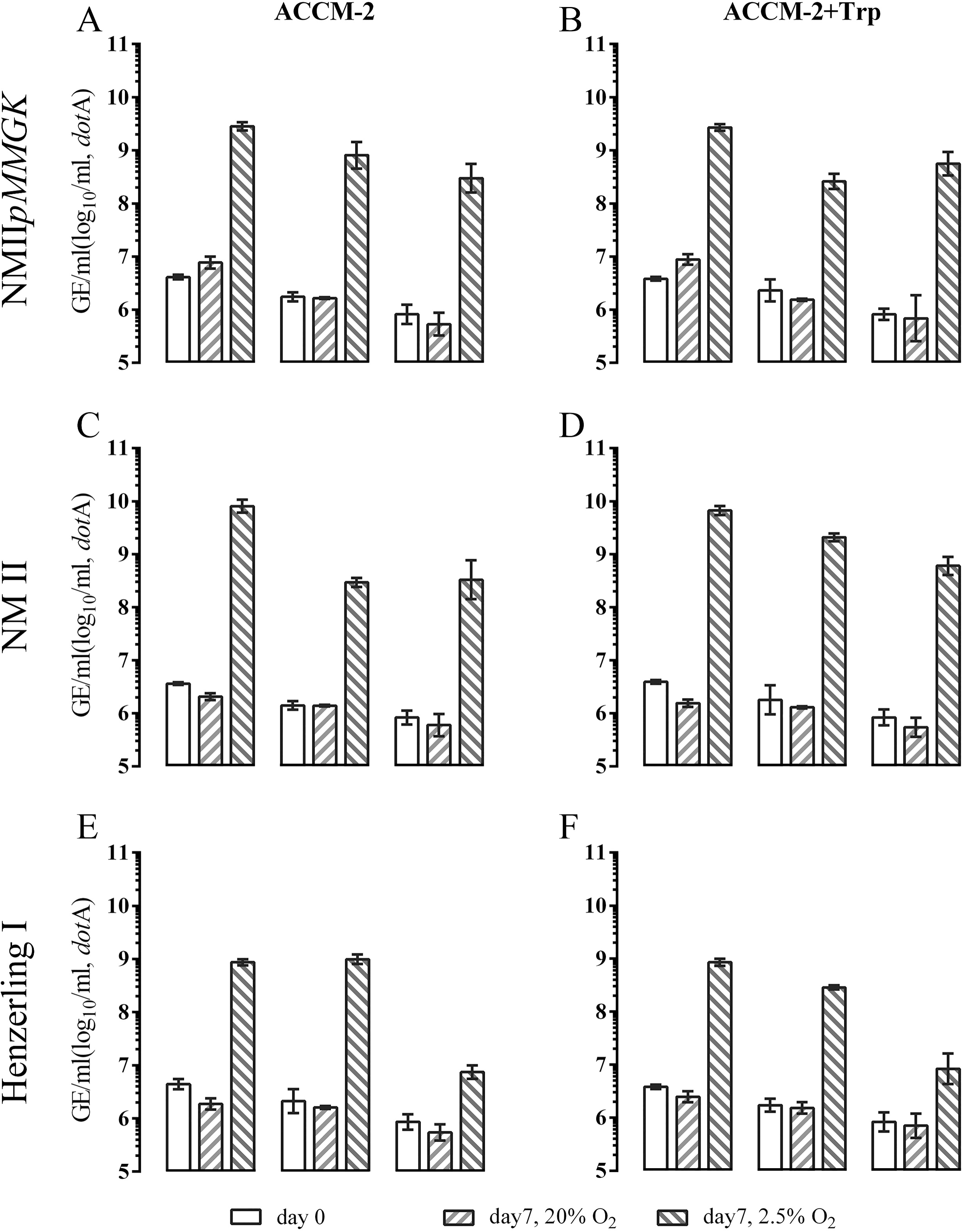
Growth of frozen stocks of *C. burnetii* strains-Henzerling phase I and Nine Mile phase II and NMII*pMMGK* in ACCM-2 with **(A, C, E)** or without tryptophan **(B, D, F)** at 2.5% and 20% oxygen. In both ACCM-2 and ACCM-2 plus tryptophan, frozen stocks of three *C. burnetii* strains have significant growth at 2.5% oxygen. However, frozen stocks of all strains fail to grow at 20% oxygen, regardless of the media and their inoculum concentrations.

### Phase II *C. burnetii* transformants show enhanced virulence in the SCID mouse infection model

The SCID mouse model allows for assessing *in vivo* pathogenesis of phase II *C. burnetii* (17). Our *C. burnetii* phase II strains with or without the pMMGK plasmid showed variable growth capability under normoxia. We next intended to investigate whether pMMGK affects the virulence of phase II *C. burnetii* in the SCID mouse model. Firstly, fresh (one day old) inocula of 1 × 10^8^ genome equivalents (GE) of NMII or NMII*pMMGK* were used to infect four weeks old female SCID mice by peritoneal (IP) injections. 16 days post infection similar magnitude of splenomegaly was detected in both infected groups (*p*=0.10) (Figure 4A, C). Notably, two cases of exceptional splenomegaly were observed in the NMII*pMMGK* group. To examine the potential influences of inoculum age on *in vivo* virulence, another infection experiment using frozen bacterial stocks and five weeks old female SCID mice were conducted. Compared to the results of using one day old inocula, 18 days post infection one-month-old frozen stock of NMII caused similar size of splenomegaly (Figure 4B, C). Unexpectedly, however, frozen stocks of both one month old and 19 months old of NMII*pMMGK* induced similar large splenomegaly in all infected mice (Figure 4B, C). The sizes of splenomegaly are in accordance with genome equivalents in all infected mice (Figure 4D). Taken together, frozen stocks of phase II *C. burnetii* transformants showed enhanced virulence in the SCID mouse model.

**Figure 4.**
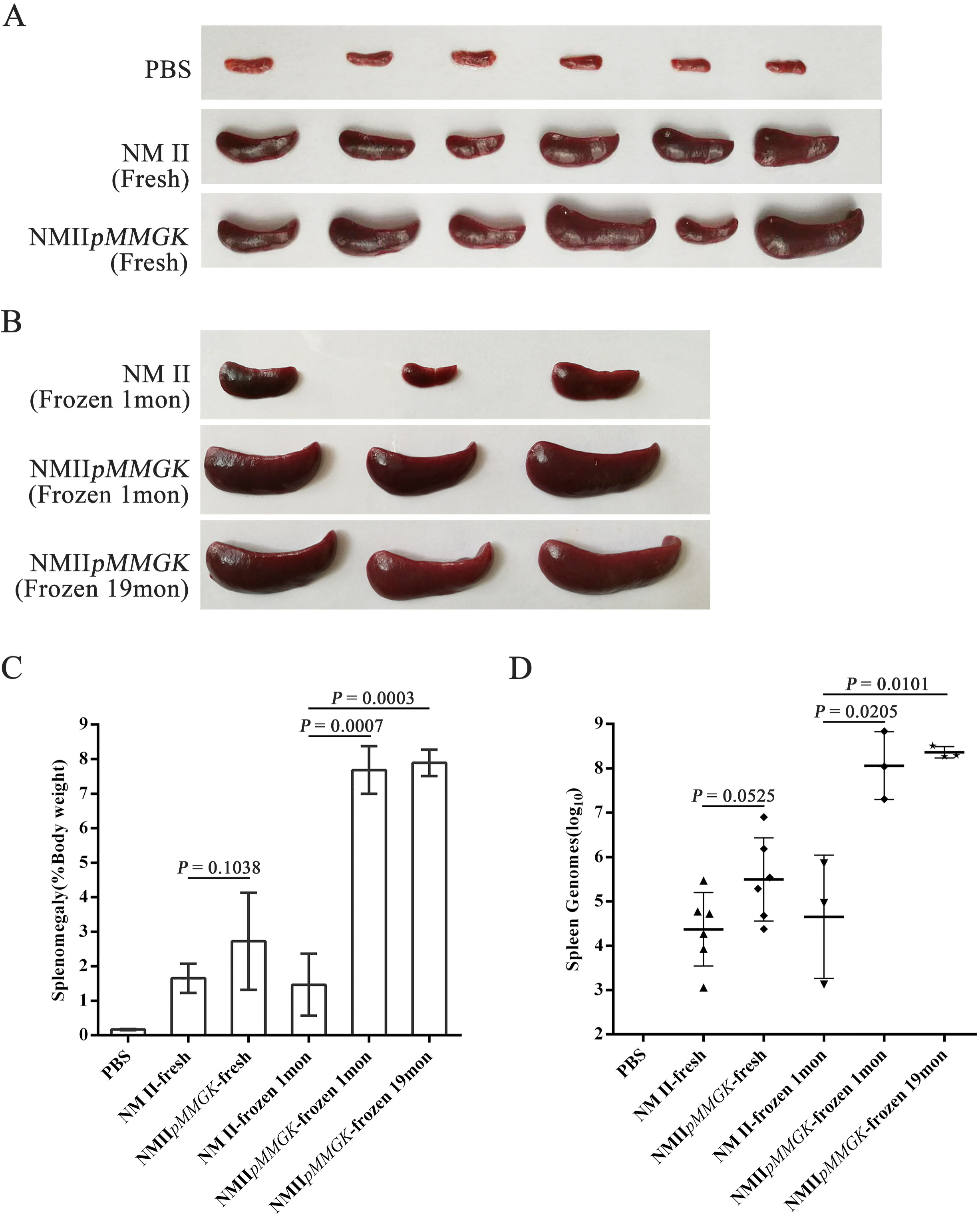
Intra-peritoneal challenge of SCID mice with different inoculum ages of *C. burnetii* strains-Nine Mile phase II and NMII*pMMGK*. **(A, B)** Spleens were removed from challenged and control mice. **(C)** Splenomegaly calculated as spleen weight as a percentage of total body weight at the time of necropsy after infection. **(D)** Genome equivalents calculated using Taqman real-time PCR with DNA purified from infected lungs. Error bars represent standard deviation from the mean.

## Discussion

The development of the axenic culture medium significantly advanced *C. burnetii* research (11). The axenic culture medium for *C. burnetii* was continuously improved (11-14). These medium studies found that hypoxic conditions are strictly required for *C. burnetii* axenic replication. In this study, we characterized the axenic growth of different *C. burnetii* strains in ACCM-2 with or without tryptophan supplementation, especially their growth under normoxic conditions. Our goal was to further improve the axenic culture of *C. burnetii* under normoxia.

We found that, under normoxic conditions if starting from appropriate concentrations of fresh inoculum, *C. burnetii* transformants of pMMGK can have robust growth in ACCM-2, regardless of tryptophan supplementation. Similarly, wild type *C. burnetii* phase II can grow significantly in ACCM-2 with tryptophan supplementation. Clearly, the *C. burnetii* transformants have a greater capability of growing under normoxia. The mechanism of *C. burnetii* normoxic growth is unknown, but likely its expression of *cyoABCDE* and *cydAB* is coordinately regulated in a reciprocal fashion in response to oxygen concentration as in *E. coli* (23).

*C. burnetii* has been recognized as an intracellular parasite in nature. Inside the infected cells, *C. burnetii* replicates in a hypoxic, phagolysosome-like vacuole (24). The cytochrome *bd* oxidase, with its increased affinity for oxygen, is likely used by *C. burnetii* under hypoxic conditions. The alternative terminal ubiquinol oxidase in *C. burnetii* -cytochrome *o* (typically used for aerobic respiration), correlates with our finding of normoxic growth. Intriguingly, the normoxic growth is affected by multiple factors including the shuttle vector pMMGK, the age and concentration of inoculum, and the phase of *C. burnetii*. The differential normoxic growth of different strains suggests that, in *C. burnetii* phase II, a temperature-sensitive quorum sensing signal molecule like RNA might be involved in the regulated expression of genes for aerobic respiration, especially cytochrome *o*.

The capability of normoxic growth in ACCM-2 without tryptophan supplementation is restricted to *C. burnetii* transformants. Their enhanced virulence is likely associated with normoxic growth. How might the shuttle vector pMMGK confers these “gain-of-function” phenotype to *C. burnetii*? The shuttle vector pMMGK brings new proteins (RepA, RepB, RepC, eGFP and Kan^R^) and new RNAs to *C. burnetii*. The potential effects of these proteins and RNAs on bacterial physiology are impossible to predict (25, 26). The RSF1010 *ori*-based shuttle vectors are commonly used for *C. burnetii* transformation (27, 28). Whether pMMGK or other similar vectors can restore normoxic growth of other *C. burnetii* phase I and phase II strains will be worthy of further investigation.

The normoxic cultures in this study were performed under static conditions. Given that oxygen supply in static cultures is much less than in shaken cultures, it is necessary to test *C. burnetii* propagation in normoxic shaken cultures. This will help determine the influence of inoculum concentration on *C. burnetii* growth under normoxia, as one may argue that high concentrations of inoculum might reduce the physical concentration of oxygen in media. Despite this possibility, static cultures of *C. burnetii* under normoxic and hypoxic conditions have distinct outcomes in infected macrophages (29). *C. burnetii* replication in macrophages is prevented in static hypoxic cultures, but not in static normoxic cultures (29).

In conclusion, we characterized the growth of various *C. burnetii* strains in ACCM-2 with or without tryptophan under normoxic and hypoxic conditions. Consistent with previous reports, *C. burnetii* had robust growth under hypoxia. What is surprising is that we found a *C. burnetii* phase II transformant and the wild type *C. burnetii* phase II can have significant to robust growth under certain normoxic circumstances. Compared to the wild type strain, the transformant displayed enhanced virulence in the SCID mouse model. Normoxic growth of *C. burnetii* is affected by the shuttle vector pMMGK, the age and concentration of inoculum, and the phase of *C. burnetii*. Our findings suggest that current axenic culture of *C. burnetii* under hypoxia might possibly be further modified to allow routine axenic culture under normoxia.

## Acknowledgements

This work was supported by National Natural Science Foundation of China (No. 31570177), Fundamental Research Funds for Central Universities (No. BUCTRC201917), Key Project of Beijing University of Chemical Technology (No. XK1803-06) and Foundation of State Key Laboratory of Pathogen and Biosecurity (No. SKLPBS1409).

